# VueGen: Automating the generation of scientific reports

**DOI:** 10.1101/2025.03.05.641152

**Authors:** Sebastian Ayala-Ruano, Henry Webel, Alberto Santos

## Abstract

**Motivation:** The analysis of omics data typically involves multiple bioinformatics tools and methods, each producing distinct output files. However, compiling these results into comprehensive reports often requires additional effort and technical skills. This creates a barrier for non-bioinformaticians, limiting their ability to produce reports from their findings. Moreover, the lack of streamlined reporting workflows impacts reproducibility and transparency, making it difficult to communicate results and track analytical processes.

**Results:** We present VueGen, a tool that automates the creation of reports from bioinformatics outputs, allowing researchers with minimal coding experience to communicate their results effectively. With VueGen, users can produce reports by simply specifying a directory containing output files, such as plots, tables, networks, Markdown text, and HTML components, along with the report format. Supported formats include documents (PDF, HTML, DOCX, ODT), presentations (PPTX, Reveal.js), Jupyter notebooks, and Streamlit web applications. To showcase VueGen’s functionality, we present two case studies and provide detailed documentation to help users generate customized reports.

**Availability and implementation:** VueGen is distributed as a Python package, Docker image, nf-core module, and desktop application. The source code is freely available on https://github.com/Multiomics-Analytics-Group/vuegen under the MIT license. Documentation is provided at https://vuegen.readthedocs.io/.

## 1 Introduction

The analysis of omics data and other biological research produces many outputs, including plots, tables, descriptions, statistics, and other data. As the results and analyses grow, organizing these outputs in a structured format becomes increasingly challenging (Kreiman and Maunsell, 2011). Moreover, aggregating and sharing these results can be difficult because of the lack of standardization, slowing down the collaboration process (Borgman, 2012).

Reports are crucial in addressing the challenges of organizing and communicating research findings. A structured report provides a comprehensive summary of the project’s outputs and ensures consistency in the presentation of data. It also facilitates sharing results with collaborators and promotes research accountability and reproducibility (Macleod *et al*., 2021; Julpisit and Esichaikul, 2019).

However, generating and formatting scientific reports from multiple output files and making them accessible can be complex, especially for researchers without a technical background. The process often requires coding skills or familiarity with specialized software, making it time-consuming and error-prone, particularly when handling large datasets and outputs. Furthermore, non-technical users may struggle to organize results consistently, leading to errors (Kumuthini *et al*., 2020).

There are various reporting tools such as Quarto, Tableau, PowerBI, and MultiQC, each with distinct strengths and limitations. Quarto (Allaire *et al*., 2024) requires basic technical expertise and does not support dynamic web applications by default. Tableau (Beard and Aghassibake, 2021) and PowerBI (https://www.microsoft.com/en-us/power-platform/products/power-bi) are powerful for data visualization and business intelligence, but these tools rely heavily on graphical interfaces, require significant manual work to produce consistent reports, offer restricted features on their free versions, and are not designed to integrate different data types. MultiQC (Ewels *et al*., 2016), while the *de facto* tool for generating quality control reports, is limited to HTML reports, offering no flexibility in output formats and data types. Moreover, adding new quality control metrics and features requires extending the core MultiQC, which can be challenging for non-technical users. VueGen addresses these limitations by automating the creation of reports in multiple formats from diverse bioinformatics outputs. With VueGen, users can produce reports by simply specifying a directory containing output files, such as plots, tables, networks, Markdown text, and HTML components, along with the report format. Supported formats include documents (PDF, HTML, DOCX, ODT), presentations (PPTX, Reveal.js), Jupyter notebooks, and Streamlit web applications. **Figure 1** presents a graphical overview of the software’s workflow and components. With a modular and cross-platform design, this software facilitates integration into various analytical workflows and enables researchers with minimal or no coding experience to communicate their results effectively. As a benefit, the focus shifts from time-consuming report generation to the meaningful analysis and interpretation of results.

**Figure 1.**
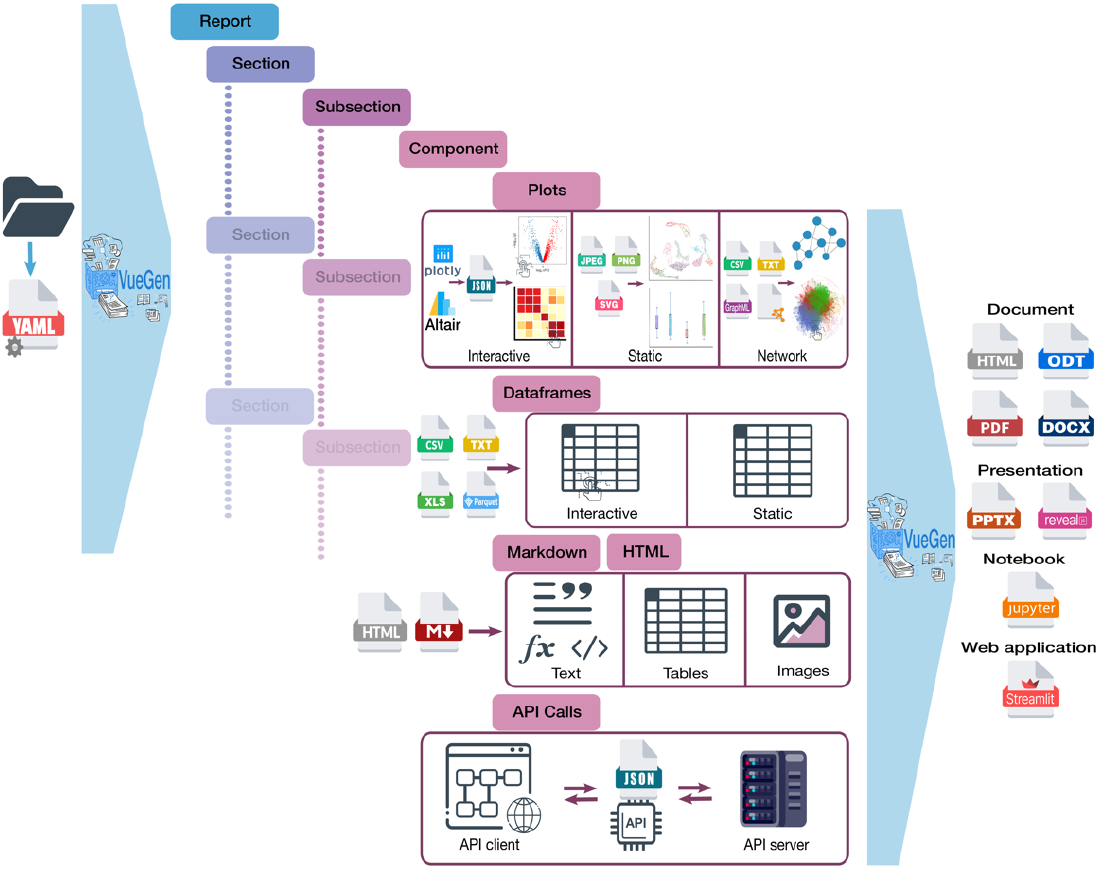
Graphical overview of VueGen’s workflow and components. The diagram illustrates VueGen’s pipeline and report structure, starting from either a directory or a YAML configuration file. If a directory is provided, the software generates a configuration file with the report structure. Reports consist of hierarchical sections and subsections, each containing various components such as plots, dataframes, Markdown, HTML, and data retrieved via API calls. Depending on the file format and report type, components can be static or interactive. Reports can be exported in various formats, including documents (PDF, HTML, DOCX, ODT), presentations (PPTX, Reveal.js), Jupyter notebooks, and Streamlit web applications.

## 2 Implementation

VueGen offers various implementation options for both non-technical and experienced users. It is available as a Python package, Docker image, nf-core module, and cross-platform desktop application with a user-friendly interface, making it accessible and customizable for different user needs and expertise levels. To streamline software maintenance, VueGen applies a continuous integration and deployment (CI/CD) pipeline for automated releases of the tool and its documentation.

### 2.1. Python Package

VueGen is an open-source tool developed in Python (https://github.com/Multiomics-Analytics-Group/vuegen), using various tools from Python’s standard library and its data science ecosystem and supporting versions 3.9 to 3.13. The package adheres to official Python standards and is available on the Python Package Index (PyPI) and Bioconda (Grüning *et al*., 2018). To simplify the installation process and ensure reproducibility, a Docker image is also available at https://quay.io/repository/dtu_biosustain_dsp/vuegen.

As shown in **Figure 2A**, the software architecture consists of three main parts: (i) managing input files and metadata that compose the report content, obtained from either a configuration file or a directory, (ii) defining the report structure using classes that represent sections, subsections, and components (e.g., plots, dataframes, markdown files, and API calls), and (iii) creating reports through classes that handle the logic for various dynamic and static report types, including Streamlit, HTML, PDF, DOCX, ODT, Reveal.js, PPTX, and Jupyter. **Figure S1** presents an extended class diagram with attributes and methods. VueGen’s modules and functionality are well documented at https://vuegen.readthedocs.io/. If a directory is provided, a YAML configuration file is automatically generated from the folder structure, where first-level folders are treated as sections and second-level folders as subsections, containing the components. The titles for sections, subsections, and components are extracted from the corresponding folder and file names, and afterward, users can add descriptions, captions, and other details to the configuration file. Component types are inferred from the file extensions and names. The order of sections, subsections, and components can be defined using numerical suffixes in folder and file names. Users can customize the report by modifying the generated configuration file or creating their own based on the example files from the case studies. If a configuration file is given, users can specify titles and descriptions for sections and subsections, as well as component paths and required attributes, such as file format and delimiter for dataframes, plot types, and other details. The documentation provides configuration file examples for basic and advanced reports.

**Figure 2.**
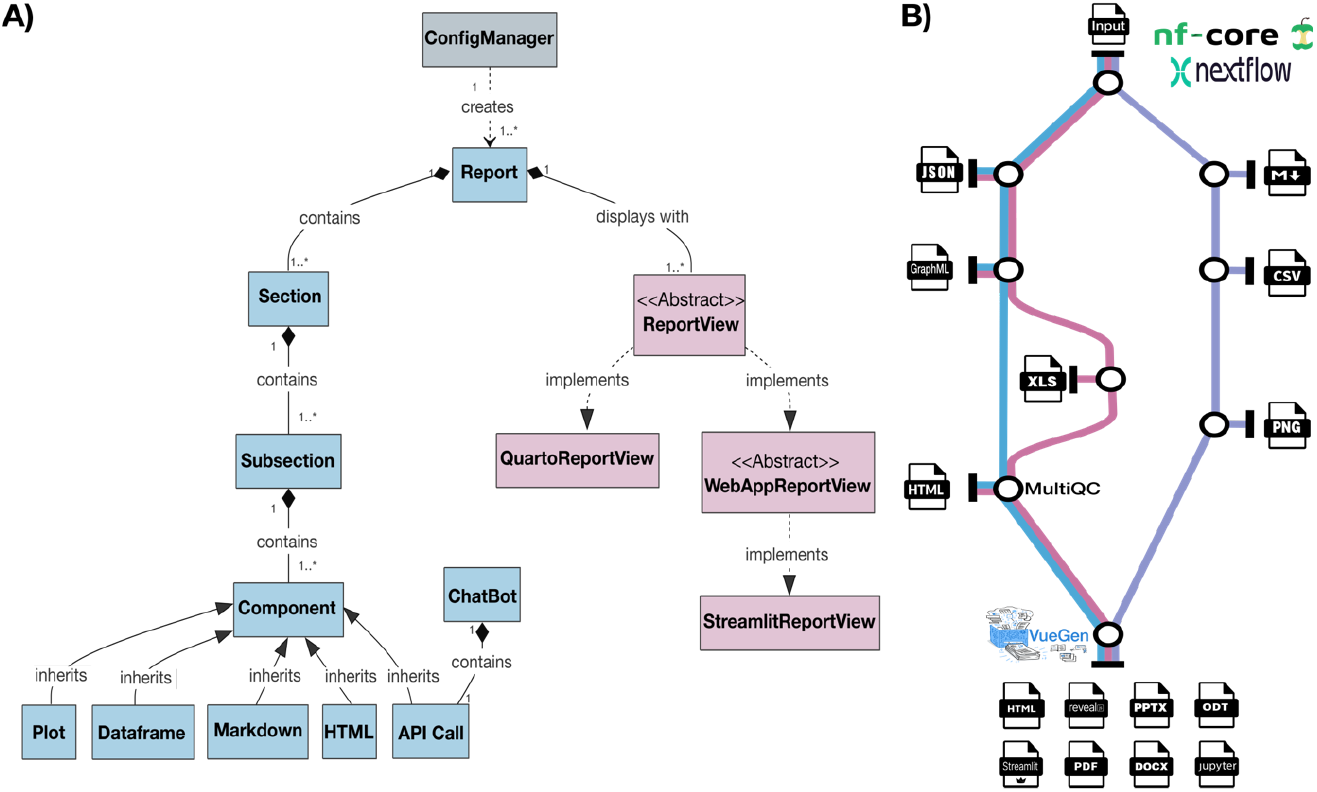
**A) VueGen class diagram.** Boxes represent classes, and arrows show their relationships. Colors indicate functionality: gray for classes managing input files and metadata, blue for those defining the report structure, and red for classes generating reports. **B) Metro-map representation of the VueGen nf-core module**. Stations (circles) represent processes within a module, subworkflow, or pipeline, which produces various output files. The workflow begins with an input dataset, proceeds through sequential stages that produce intermediate outputs in multiple formats, and concludes with report generation.

VueGen mainly relies on two frameworks: Streamlit (https://streamlit.io/) for creating web applications, and Quarto (Allaire *et al*., 2024) for generating the other report types. The software operates by instantiating section, subsection, and component classes from the configuration file, while a report view class manages the report structure and organization. Streamlit reports generate a directory with subfolders for every section, each containing Python scripts for individual subsections that correspond to the pages in the web application. Other report types generate a directory with a Quarto Markdown (qmd) file that structures the entire report. Once the reports are obtained, they are rendered using Streamlit or Quarto. If the report type is a Streamlit web interface, it can launch the application automatically by setting the *st_autorun* argument to True (default is False).

VueGen supports interactive plots generated with either Plotly (Plotly Technologies Inc., 2015) or Altair (Satyanarayan *et al*., 2017), provided as JSON files. It also handles static plots in PNG, JPG, SVG, and other image formats. Networks are loaded as interactive plots using Pyvis (Perrone *et al*., 2020) and can be provided as CSV or TXT files containing edge lists or adjacency matrices, or as GraphML, GML, GEXF, CYJS, or HTML files. If interactive plots are included and the report type is static (e.g., DOCX, PDF, PPTX, etc.), plots and networks are converted into static images.

Tables can be included in CSV, TXT, Parquet, and XLSX formats. Markdown and HTML files are also supported, allowing users to specify formatted text, images, tables, mathematical equations, and other content. Finally, API call components can be created by manually defining the API URL, request body, and other details in the configuration file. These components are applicable only for Streamlit reports because this framework enables sending API requests and fetching responses for real-time display, which can be very useful for retrieving additional information or annotations.

### 2.2. Nf-core Module

The VueGen nf-core module is designed to automate report generation from outputs produced by other modules, subworkflows, or pipelines. The module integrates the VueGen Python library and customizes it for compatibility with the Nextflow environment (Di Tommaso *et al*., 2017). It was built using the nf-core module template and applying the community standards (Ewels *et al*., 2020). This module is compatible and extends existing Nextflow reporting tools such as MultiQC (Ewels *et al*., 2016), AssemblyQC (Rashid *et al*., 2024), and others, integrating their HTML outputs as components within VueGen reports.

### 2.3. Article short title

**Figure 2B** shows a metro-map representation of the VueGen nf-core module. The workflow represents a traditional Nextflow workflow with an initial input dataset and multiple intermediate outputs in various formats generated in different processes. In the final stage, the VueGen nf-core module collects these outputs from a directory and generates reports in multiple formats. By simplifying and standardizing report generation, VueGen improves the accessibility and sharing of results within the Nextflow ecosystem. Integrating VueGen into nf-core pipelines will streamline reporting workflows, and provide consistent, customizable outputs across diverse omics analyses. This facilitates communication, comparison, and reproducibility of results, fostering collaboration and workflow transparency. The VueGen nf-core module code can be found at https://github.com/Multiomics-Analytics-Group/nf-vuegen.

### 2.3. Desktop application

The VueGen Python package was wrapped into a cross-platform desktop application compatible with macOS and Windows. This makes the tool accessible to users without coding experience, allowing them to interact with it through an intuitive graphical interface, without requiring command-line usage or the installation of Python or Nextflow. It uses Tkinter, a modern layout for graphical user interfaces (GUI), bundled as a self-contained application using PyInstaller. The executables files for installing the GUI are available on the GitHub release page: https://github.com/Multiomics-Analytics-Group/vuegen/releases.

## 3 Case Studies

Two case studies were designed to demonstrate Vuegen’s functionality. The first is a basic example that uses a predefined directory, while the second is an end-to-end example that generates the directory and components using data from the Earth Microbiome Project (EMP) (Thompson *et al*., 2017). Both case studies were implemented with the VueGen Python package in Jupyter notebooks and the nf-core module in Nextflow scripts.

### 3.1. Predefined Directory

This introductory case study familiarizes users with the tool, its report types, file formats, and other features. In this example, a directory with example plots, dataframes, Markdown, and HTML components was created. The notebook and Nextflow script load this directory and generate reports in different formats. After generating a report from the directory, the configuration file can be modified to add new sections, subsections, and components. The notebook example can be executed in Google Colab using the following link: https://colab.research.google.com/github/Multiomics-Analytics-Group/vuegen/blob/main/docs/vuegen_basic_case_study.ipynb.

### 3.2. Earth Microbiome Project Data

This advanced case study demonstrates the application of VueGen in a real-world scenario, starting from content generation. The EMP is an initiative to characterize global microbial taxonomic and functional diversity (Thompson *et al*., 2017). **Figure 3A** shows example plots from the original study. EMP data, scripts, and outputs are publicly available at https://github.com/biocore/emp.

**Figure 3.**
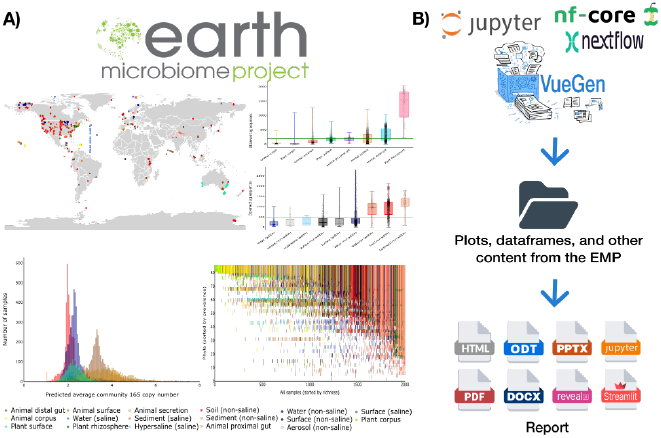
Case study: VueGen application to the Earth Microbiome Project (EMP). **A)** EMP logo and example plots from the original study, illustrating global microbial diversity. **B)** Workflow to reproduce EMP visualizations and create reports with VueGen. A Jupyter notebook and Nextflow script were used to generate plots, dataframes, descriptions, and other content. These outputs were organized into a folder and processed with VueGen to create reports, demonstrating its application in a research project.

The notebook and Nextflow script process the EMP data, create plots, dataframes, and other components, and organize outputs within a directory to produce reports (**Figure 3B**). Report content and structure can be adapted by modifying the configuration file. Each report consists of sections on exploratory data analysis, metagenomics, and network analysis. In the Nextflow script, each section corresponds to a process that generates outputs, which the nf-core module then uses to build a report. This example illustrates how the VueGen nf-core module can be integrated into a pipeline to produce reports as the final stage (**Figure 2B**). HTML and Streamlit report examples of the current VueGen release are available at https://multiomics-analytics-group.github.io/vuegen/ and https://earth-microbiome-vuegen-demo.streamlit.app/. The notebook example can be executed in Google Colab using the following link: https://colab.research.google.com/github/Multiomics-Analytics-Group/vuegen/blob/main/docs/vuegen_case_study_earth_microbiome.ipynb.

## 4 Conclusions and Future Perspectives

Organizing and sharing research outputs in a standardized format is often complex and time-consuming. VueGen addresses this challenge by automating report generation, making it accessible to researchers with minimal or no coding expertise. The software supports static and interactive data, structured in various report formats. VueGen is distributed as a Python package, Docker image, nf-core module, and cross-platform desktop application, enabling users to integrate it into their workflows according to their specific requirements. Extensive documentation of the package’s classes and functions is provided. Moreover, two case studies demonstrate its application in research projects.

Potential future developments include automating report deployment to Streamlit Cloud, GitHub pages, or other services. Further, we aim to integrating VueGen with other bioinformatics analysis and visualization tools to support entire analytical workflows. Additionally, we will expand the support for more data types and web application frameworks (e.g., Shiny, Dash, Panel), and simplify the report generation process.

## Acknowledgments

VueGen is built upon numerous open-source projects, including Python, Streamlit, Quarto, Nextflow, nf-core, and others. We acknowledge the authors for their contributions, which made this work possible.

## Conflict of interest

The authors declare no conflict of interest.

## Funding

This work has been supported by the grant number NNF20CC0035580.

## Data and code availability

VueGen is distributed as a Python package, Docker image, nf-core module, and desktop application. The VueGen Python package code, installation instructions, and case study notebooks are available at https://github.com/Multiomics-Analytics-Group/vuegen. The Docker image is available at https://quay.io/repository/dtu_biosustain_dsp/vuegen. The nf-core module code can be found at https://github.com/Multiomics-Analytics-Group/nf-vuegen. The executables files for installing the GUI are available on the GitHub release page: https://github.com/Multiomics-Analytics-Group/vuegen/releases. Documentation is accessible at https://vuegen.readthedocs.io/. HTML and Streamlit report examples of the current VueGen release are available at https://multiomics-analytics-group.github.io/vuegen and https://earth-microbiome-vuegen-demo.streamlit.app/. All code from this project is open and shared under the MIT license.

## Notes

https://github.com/Multiomics-Analytics-Group/vuegen

https://vuegen.readthedocs.io/

### Competing Interest Statement

The authors have declared no competing interest.

