## Supplementary figures and images for "VueGen: Automating the generation of scientific reports"

### Supplementary Figure 1

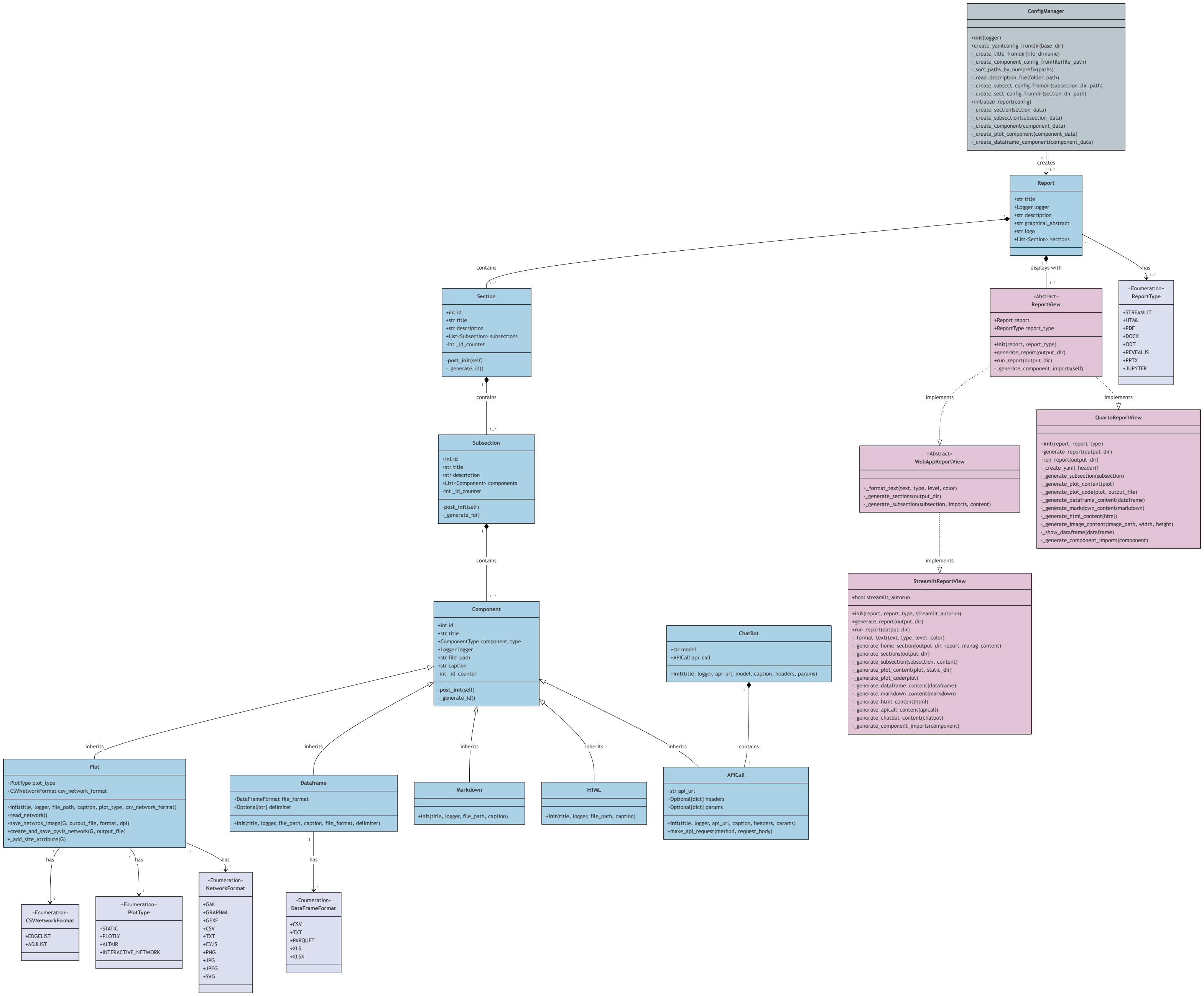
